# Mega- and meta-analyses of fecal metagenomic studies assessing response to immune checkpoint inhibitors

**DOI:** 10.1101/2021.04.27.441693

**Authors:** Alya Heirali, Bo Chen, Matthew Wong, Pierre HH Schneeberger, Victor Rey, Anna Spreafico, Wei Xu, Bryan A. Coburn

## Abstract

**Purpose:** Gut microbiota have been associated with response to immune checkpoint inhibitors (ICI) including anti-PD-1 and anti-CTLA-4 antibodies. However, inter-study difference in design, patient cohorts and data analysis pose challenges to identifying species consistently associated with response to ICI or lack thereof.

**Experimental Design:** We uniformly processed and analyzed data from three studies of microbial metagenomes in cancer immunotherapy response (four distinct data sets) to identify species consistently associated with response or non-response (n=190 patient samples). Metagenomic data were processed and analyzed using Metaphlan v2.0. Meta- and mega-analyses were performed using a two-part modelling approach of species present in at least 20% of samples to account for both prevalence and relative abundance differences between responders/non-responders.

**Results:** Meta- and mega-analyses identified five species that were concordantly significantly different between responders and non-responders. Amongst them, *Bacteroides thetaiotaomicron* and *Clostridium bolteae* relative abundance (RA) were independently predictive of non-response to immunotherapy when data sets were combined and analyzed using mega-analyses (AUC 0.59 95% CI 0.51-0.68 and AUC 0.61 95% CI 0.52-0.69, respectively).

**Conclusions:** Meta- and mega-analysis of published metagenomic studies identified bacterial species both positively and negatively associated with immunotherapy responsiveness across four published cohorts.

## Introduction

A number of studies have demonstrated the gut microbiome is associated with response to immune checkpoint inhibitors (ICI)^1–5^. Anti-programmed cell death protein 1(PD-1) and anti-cytotoxic T lymphocyte-associated protein 4 (CTLA-4) targeting agents derepress anti-tumor T-cells. In the past five years, mouse models and human observational cohort studies have shown that the gut microbiome of responders to ICI is compositionally different from non-responders ^1–6^. However, inter-study differences in taxa associated with response to ICIs make it challenging to discern which organisms are consistently associated with response or non-response across studies and different cancers. This variability may in part be due to differences in sequence analytical pipelines.

We conducted both meta- and mega-analyses of metagenomic studies of the gut microbiome in ICI recipients to determine species consistently enriched/depleted in responders compared to non-responders. Here, metagenomic data were selected to maximize the taxonomic resolution and delineate species-specific associations that may not be evident when taxa are annotated at the genus level.

## Methods

### Cohort Inclusion

We included three studies with publicly available metagenomic data and meta-data. Data was further divided into distinct data sets for the Routy *et al.* study to assess differences based on tumor type. These data sets include individuals with melanoma, renal cell carcinoma (RCC) and non-small cell lung cancer (NSCLC).

### Definitions

Patients were classified as responders (R) or non-responders (NR) using the response evaluation criteria in solid tumors across all three studies (RECIST v1.1)^3–5,7^. Data sets were analyzed in aggregate (meta/mega analysis) and separately. The primary outcome of interest was to detect species consistently enriched/depleted in responders to ICI across data sets. This was achieved using a number of statistical methods described below. The secondary outcome was to identify predictors of response using receiver operator characteristics area under the curve (ROCAUC) analyses.

### Microbiome Data processing

Raw sequencing data were obtained, and quality filtered using FastQC and MultiQC^8,9^. Metaphlan v2.0 was used for its fast and robust species level annotation of microbial genomes^10^. Alpha-diversity measures were calculated using Phyloseq^11^. Statistical analyses and data visualization were conducted in R and Prism^12,13^.

### Statistical Analyses

#### Two-part log-normal model

The raw sequencing data was observed as relative abundance (RA) of each species. Our target was to estimate the R/NR group differences of each species, where the differences consist of both species prevalence (presence/absence) and magnitude of abundance when species are present. We conducted two-part model analysis for species prevalence and relative abundance separately. Specifically, we used a logistic regression model to detect the association between species prevalence and R/NR. For RA data greater than zero, we observed log-normal distributions, and thus used linear regression model to find its association to R/NR group. The logistic regression estimates (effect size) represent the log odds ratio of species prevalence between R and NR group, and linear regression estimates (effect size) represent the change of mean RA between R and NR group after excluding all zeros. We calculated 95% confidence intervals (CIs) for all estimates and tested if each estimate is significantly different from 0. A positive estimate means that R had a higher proportion of non-zeros among all samples, or R had a higher RA on average compared to NR. We noted that for each species, estimates for prevalence and abundance may have different directions, which is described as “dissonant” effect ^14^. In order not to let the opposite estimates reduce statistical test power, we tested each estimate by chi-squared test without considering its direction. We then combined these two test statistics to achieve the overall p-value, which tests the overall difference between R and NR for each species.

#### Meta-analysis

Meta-analysis is a two-stage approach^15^. In the initial stage data are analyzed separately for each study. In the second-stage test results for each study are aggregated to obtain summary data across all studies^15^. On the other hand, mega-analysis is a one-stage approach that pools and analyzes raw data from a number of studies to estimate the overall effect^15^. Both approaches have their strengths and limitations^15^. We conducted a meta-analysis by combining the test results for each of the four data sets in Table 1. Logistic regression and linear regression estimates are only accurate if the zero proportion in each species is sufficiently low. Accordingly, species were excluded if their prevalence was lower than 20%. As species prevalence differs by study, some species were filtered out only in a subset of the data sets. Assuming the independence of each data set, the overall meta-analysis p-values were computed for the combined test statistic, which is the sum of the test statistics from the four data sets.

**Table 1:** Characteristics of cohorts included in meta-analysis and corresponding taxa associated with response to ICI using metagenomic analyses.

| Patient population | N (= total number of patients) | Responders | non-Responders | Organisms enriched in RECIST responders | Organism Enriched in RECIST non-responders | Reference |
| --- | --- | --- | --- | --- | --- | --- |
| Metastatic melanoma | N=25 | N=14 | N=11 | *Faecalibacterium spp.* | *Bacteroides thetaiotaomicron*,<br>*Escherichia coli*,<br>*Anaerotruncus colihominis* | Gopalakrishnan *et al.* 2018 |
| Metastatic melanoma | N=38 | N=14 | N=24 | *Enterococcus faecalis*, *E. coli*,<br>*Escherichia spp.*, *Bacteroides ovatus*,<br>*Turicibacter sanguinis*, *Clostridium nexile*, *Enterococcus faecium*,<br>*Collinsella aerofaciens*,<br>*Bifidobacterium adolescentis*,<br>*Klebsiella oxytoca*, *Veillonella parvula*,<br>*Parabacteroides merdae*,<br>*Lactobacillus* species,<br>and *Bifidobacterium longum*,<br>*Campylobacter gracilis* | *Burkholderiales bacterium 1147*,<br>*Holdemania filiformis*,<br>*Coprococcus comes* | Matson *et al.* 2018 |
| Advanced NSCLC | N=65 | N=33 | N=32 | *Akkermansia muciniphila*,<br>*Ruminococcaceae*, *Faecalibacterium*,<br>*Alistipes spp.*, *Eubacterium* species,<br>*Firmicutes*, *Cloacibacilius porcorum*,<br>*Enterococcus faecium*, *Clostridiales*,<br>*Firmicutes*, *Prevotella*, | *Prevotella spp.*,<br>*Clostridiales*,<br>*Blautia*, *Bacteroides calrus*,<br>*Proteobacteria*,<br>*Bacteroides nordii*,<br>*Parabacteroides distasonis* | Routy *et al.* 2018 |
| RCC | N=62 | N=42 | N=20 |  |  |  |

#### Mega-analysis

For each species included in the meta-analysis, we also conducted a mega-analysis. The mega-analysis simply combined all samples from all four data sets, and two-part log-normal models were implemented similarly. We also calculated the CIs for all estimates and overall chi-squared test p-values combining species prevalence and abundance, which were then compared with the meta-analysis results.

## Results

### Characteristics of cohorts included in analyses

In this mega- and meta-analysis, we included three studies that assessed fecal metagenomics of R and NR to ICI in patients with varying tumor types (Table 1). Data from Routy *et al*., was analyzed as subsets based on tumor type RCC vs advanced NSCLC for a total of 4 analyzed cohorts. DNA extraction, sequencing platform and sequencing depth varied by study and are summarized in Table S1. A total of 190 patients (n=103 R; n=87 NR) were included from the three studies.

A total of 469 species were identified across all four data sets of which 167 species met our inclusion criteria of being present in >20% of samples and were included in the two-part log-normal analysis (Table S2). A total of 34 species were differentially abundant based on response in at least one dataset or mega- and meta-analyses (Table S2).

### Species consistently enriched or depleted in responders compared to non-responders across data sets

The primary goal for our study was to determine species consistently enriched or depleted in R compared to NR across data sets. While each data set had unique signatures of differentially abundant species between R/NR, no species were identified that were statistically significantly associated with response across all four data sets. A number of species were significantly differentially enriched/depleted in R/NR per data set; Matson et al., (n=7 species), Routy et al., (RCC) (n=6), Routy et al., (NSCLC) (n=10), Gopalakrishnan et al., (n=9) (Figures S1 and S2 and Table S2). A total of three species were significantly enriched or depleted in R in two data sets: *Clostridiaceae bacterium JC118* was enriched in R in the Routy et al., (NSCLC) and Matson et al., data sets; *Bacteroides thetaiotaomicron* RA was significantly depleted in R in both the Routy et al., (RCC) and Gopalakrishnan et al., data sets; and an increase in *Lachnospiraceae bacterium 5163FAA* abundance was associated with response in both the Gopalakrishnan et al., and Routy et al., (NSCLC) data sets. While not significantly different, *Streptococcus australis* trended towards a higher relative abundance in R across all cohorts. In contrast, *B. thetaiotaomicron* trended towards lower relative abundance in R across all data sets. In addition, we were interested in assessing species that are consistently enriched or depleted in R across data sets based on prevalence (Figure S2 and Table S2). *Clostridium bolteae, Escherichia coli*, *Flavonifractor plautii*, *Ruminococcus lactaris* and *Streptococcus australis* were consistently less prevalent in R. In contrast, *Bacteroides caccae*, *Barnesiella intestinihominis* and *Lachnospiraceae bacterium 8157FAA* were more prevalent in R. Only one species had statistically significant but opposing associations between cohorts; *Ruminococcus gnavus* was more abundant in R in the Routy et al., (RCC) cohort and less abundant in the cohort in the Gopalakrishnan et al., study.

### Meta-analysis reveals species consistently associated with response to immunotherapy

A summary of a metagenomic meta-analysis of species level ICI response associations is shown in Figure 1. A total of thirteen species were significantly differentially abundant between R and NR in the meta-analysis. Of these thirteen species, twelve were identified as significantly different between R and NR in at least one data set on its own. *Clostridium hathewayi* was the only species identified uniquely in the meta-analysis. Five species which were significantly associated with R/NR in one dataset demonstrated a non-significant trend in the meta-analysis: *Coprobacillus spp*, *Parabacteroides spp.*, Lachnospiraceae bacterium 8157FAA, *Eubacterium ramulus*, and *Clostridium symbiosum*. Lastly, 16 species were not statistically significantly different nor trended towards response using the meta-analysis despite being significantly different in at least one dataset.

**Figure 1:**
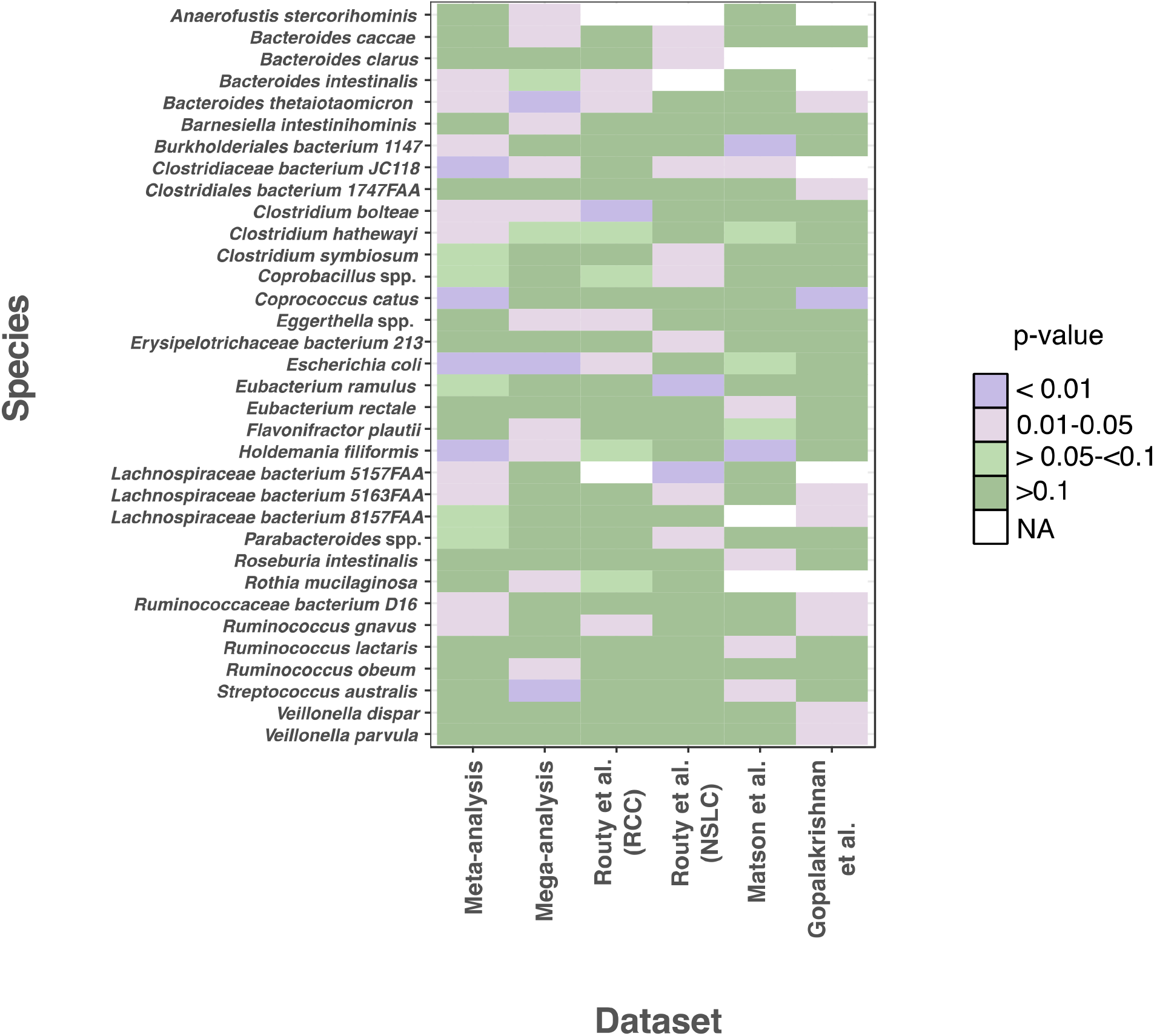
Taxa that are significantly different between responders and non-responders in at least one dataset and or mega/meta-analyses using a two-part log-normal analysis accounting for both taxa abundance and prevalence.

### Mega-analysis assessing differences in responders vs non-responders across all data sets

Mega-analysis allowed us to determine the effect size of the differences between R and NR for both prevalence and relative abundance of species across all studies (Figures S1 and S2, and Table S2). In the mega-analysis, thirteen species were statistically differentially abundant between R and NR (Figure 1). Of these thirteen species, eight were associated with response in at least one dataset. Five species were uniquely associated with response using the mega-analysis including *Anaerofustis stercorihominis, Flavonifractor plautti, Ruminococcus obeum, Rothia mucilaginosa* and *Barnesiella intestinitominis*. Of the 34 species associated with response in at least one data set or the meta-analysis, 19 species were not associated with response in the mega-analysis.

### Identification of species consistently associated with response in meta- and mega-analyses and analysis of sensitivity and specificity to predict response

We next identified species that were concordantly associated with response in both the meta and mega-analyses. Using this criterion five species were identified including *B. thetaiotaomicron*, *Clostridium bolteae, H. filiformis, Clostridiaceae bacterium JC118* and *E. coli*. We tested the sensitivity and specificity of the relative abundance of these five organisms and alpha-diversity measures to predict response to ICI using a ROCAUC analyses (Table 2). Our findings demonstrate inter-study differences in the ability to discriminate R vs NR based on species RA. No species or alpha-diversity metric consistently demonstrated a significant AUC across studies. However, each dataset had at least two species or alpha-diversity measures with AUC’s >0.60. In the Matson et al., dataset *H. filiformis* RA significantly discriminated between R/NR with an AUC of 0.72 [95% CI 0.55-0.88]. *C. bolteae* RA had a significant AUC of 0.73 [95% CI 0.60-0.87] in the Routy et al., (RCC) dataset suggesting that a higher RA of this species discriminates between R and NR. In the Gopalakrishnan et al., dataset both *E. coli* and *B. thetaiotaomicron* RA had significant sensitivity and specificity to detect response. When all four cohorts were combined sensitivity and specificity of predicting response were significant for the RA of both *C. bolteae* and *B. thetaiotaomicron.* SDI did not discriminate between R/NR.

**Table 2:** Receiver operator characteristic curve area under the curve (AUC) analysis between responders and non-responders for core taxa and alpha-diversity measures.

| Study |  |  |  |  |
| --- | --- | --- | --- | --- |
| **Matson *et al.*, (n=38 samples)** | **AUC** | **SE<sup>^</sup>** | **95% CI<sup>+</sup>** | **P-value** |
| SDI<sup>†</sup> | 0.52 | 0.10 | 0.33-0.72 | 0.81 |
| SIM<sup>‡</sup> | 0.55 | 0.10 | 0.35-0.74 | 0.63 |
| *Escherichia coli* | 0.64 | 0.10 | 0.44-0.84 | 0.15 |
| *Clostridiaceae bacterium JC118* | 0.63 | 0.10 | 0.43-0.82 | 0.19 |
| *Holdemanian filiformis* | 0.72 | 0.085 | 0.55-0.88 | **0.027** |
| *Clostridium bolteae* | 0.57 | 0.097 | 0.39-0.76 | 0.45 |
| *Bacteroides thetaiotaomicron* | 0.61 | 0.092 | 0.43-0.79 | 0.25 |
| **Routy *et al.*, (RCC, n=62)** |  |  |  |  |
| SDI | 0.51 | 0.080 | 0.35-0.67 | 0.88 |
| SIM | 0.53 | 0.079 | 0.37-0.68 | 0.74 |
| *Escherichia coli* | 0.54 | 0.075 | 0.40-0.69 | 0.58 |
| *Clostridiaceae bacterium JC118* | 0.63 | 0.077 | 0.47-0.78 | 0.11 |
| *Holdemanian filiformis* | 0.53 | 0.086 | 0.36-0.69 | 0.76 |
| *Clostridium bolteae* | 0.73 | 0.070 | 0.60-0.87 | **0.0030** |
| *Bacteroides thetaiotaomicron* | 0.55 | 0.090 | 0.37-0.72 | 0.55 |
| **Routy *et al.*, (NSCLC, n=65)** |  |  |  |  |
| SDI | 0.62 | 0.071 | 0.48-0.76 | 0.096 |
| SIM | 0.61 | 0.071 | 0.47-0.75 | 0.13 |
| *Escherichia coli* | 0.56 | 0.072 | 0.42-0.71 | 0.37 |
| *Clostridiaceae bacterium JC118* | 0.58 | 0.071 | 0.44-0.72 | 0.28 |
| *Holdemanian filiformis* | 0.50 | 0.072 | 0.36-0.64 | 0.97 |
| *Clostridium bolteae* | 0.52 | 0.073 | 0.37-0.66 | 0.81 |
| *Bacteroides thetaiotaomicron* | 0.57 | 0.072 | 0.43-0.71 | 0.33 |

| **Gopalakrishnan *et al.*, (n=25)** |  |  |  |  |
| --- | --- | --- | --- | --- |
| SDI | 0.55 | 0.13 | 0.30-0.79 | 0.70 |
| SIM | 0.64 | 0.12 | 0.41-0.88 | 0.25 |
| *Escherichia coli* | 0.73 | 0.11 | 0.52-0.93 | **0.055** |
| *Clostridiaceae bacterium JC118* | 0.50 | 0.12 | 0.27-0.74 | 0.98 |
| *Holdemanian filiformis* | 0.55 | 0.13 | 0.31-0.81 | 0.64 |
| *Clostridium bolteae* | 0.63 | 0.12 | 0.40-0.86 | 0.27 |
| *Bacteroides thetaiotaomicron* | 0.76 | 0.10 | 0.55-0.96 | **0.029** |
| **Mega-analysis (n=190)** |  |  |  |  |
| SDI | 0.56 | 0.042 | 0.47-0.64 | 0.18 |
| SIM | 0.55 | 0.042 | 0.46-0.63 | 0.28 |
| *Escherichia coli* | 0.53 | 0.042 | 0.45-0.62 | 0.44 |
| *Clostridiaceae bacterium JC118* | 0.54 | 0.042 | 0.46-0.62 | 0.33 |
| *Holdemanian filiformis* | 0.54 | 0.043 | 0.46-0.63 | 0.27 |
| *Clostridium bolteae* | 0.61 | 0.041 | 0.52-0.69 | **0.013** |
| *Bacteroides thetaiotaomicron* | 0.59 | 0.043 | 0.51-0.68 | **0.027** |

† Shannon alpha-diversity index
‡ Simpson alpha-diversity index
Δ Standard Error
+ Confidence Interval

## Discussion

In our study, we leveraged the additional statistical power of mega and meta-analyses and taxonomic resolution of metagenomic data to identify species consistently associated with R or NR across three studies (four cohorts) in which gut microbial community composition has been previously associated with response to ICI ^3–5^. We conducted a two-part log-normal analysis that accounts for the skewness and non-Gaussian distribution of microbiome data^14,16–18^. Compared to simply assessing changes in relative abundance of species, this approach has the advantage of accounting for both prevalence and relative abundance data ^14,16–18^. After uniform bioinformatic analysis, no species were identified that had a significant association with response across individual studies, but five were identified in combined analysis. Importantly, these associations were identified in spite of the inter-cohort heterogeneity in tumor type, treatment regimens, geographic location and sequencing methods, suggesting that they may be more generalizable.

Our study confirms findings of some previously identified R/NR associated species^2–5,19^. Notably, a higher abundance of *Bacteroides* has been associated with lack of response whereas a higher abundance of *Firmicutes* has been associated with response to ICI. Specifically, we observed R had a higher abundance of *Clostridiaceae bacterium JC118* and *E. coli* and a lower abundance of *B. thetaiotaomicron, H. filiformis* and *C. bolteae* compared to NR. In addition, we demonstrate that of the five organisms that were concordantly associated with response in the meta and mega-analyses, a higher relative abundance of *B. thetaiotaomicron* and *C. bolteae* were predictive of NR using a ROCAUC analyses. These data suggest that strategies that deplete non-response associated microbes (such as specific antibiotics or bacteriophages) may be viable therapeutic approaches. This is in contrasts to “pro-microbial” approaches currently being investigated in many clinical trials such as the use of probiotics, fecal microbiome transplants and stool substituents to augment response to ICI ^20^.

We observed a number of discrepancies to previously published data. Of interest, *A. muciniphila*, *B. longum* and *F. prausnitzii* were associated with response in the original studies however we did not detect differences between R and NR in our analyses. While limitations in statistical power based on sample size may partly explain this, inter-cohort variability in treatment regimens and the biology of host-treatment-microbe interactions may be important additional factors. As the mechanisms conferring microbe-induced ICI-responsiveness are being elucidated, it is clear that some will be tumor-type, host or even tumor-specific, such as the molecular mimicry of tumor antigens in gut microbes^21^, which will only confer microbe-response associations in a defined number of patients with specific tumors. Other factors, such as diet-microbe-metabolite-immune interactions may also drive tumor, host or population-specific associations which do not generalize^6^.

We acknowledge a number of limitations to our study. Firstly, differences in sample collection, DNA extraction methods and sequencing platforms from each study may bias findings- making it challenging to detect universal signals associated with response to ICI. Implementing standardized methods for microbiome studies will reduce experimental and technical biases and improve our ability to detect differences between R and NR. Second, patient heterogeneity in disease stage, diet, sex, treatment, co-morbidities limit the ability of meta or mega-analytical approaches to identify associations that may exist in only defined subsets of patients. Our sample size of 190 metagenomic samples, while as large as some cohorts analyzed by lower taxonomic resolution methods such as 16S rRNA sequencing, was still limited. Additional multi-centre prospective studies are required to validate the associations identified in this analysis and their diagnostic or therapeutic utility.

## Conclusion

Our study confirms previous findings suggesting that there are differences in the gut microbiome of R vs NR to ICI. Despite consistent bioinformatic analyses no species were found to be consistently differentially abundant between R/NR across data sets. However, using meta- and mega-analyses we identified five species that were concordantly differentially abundant between R and NR. Of these five organisms, *B. thetaiotaomicron* and *C. bolteae* were predictors of NR to ICI. These data suggest future clinical trials should assess the use of narrow spectrum anti-biotics targeting NR associated species.

## Acknowledgments

The authors acknowledge and are grateful for the support of the Tomcyzk AI and Microbiome Working Group. AH was supported by the Tomcyzk AI and Microbiome Working Group and the Princess Margaret Cancer Foundation.

