## Supplementary material for "Mega- and meta-analyses of fecal metagenomic studies assessing response to immune checkpoint inhibitors": Figure S1

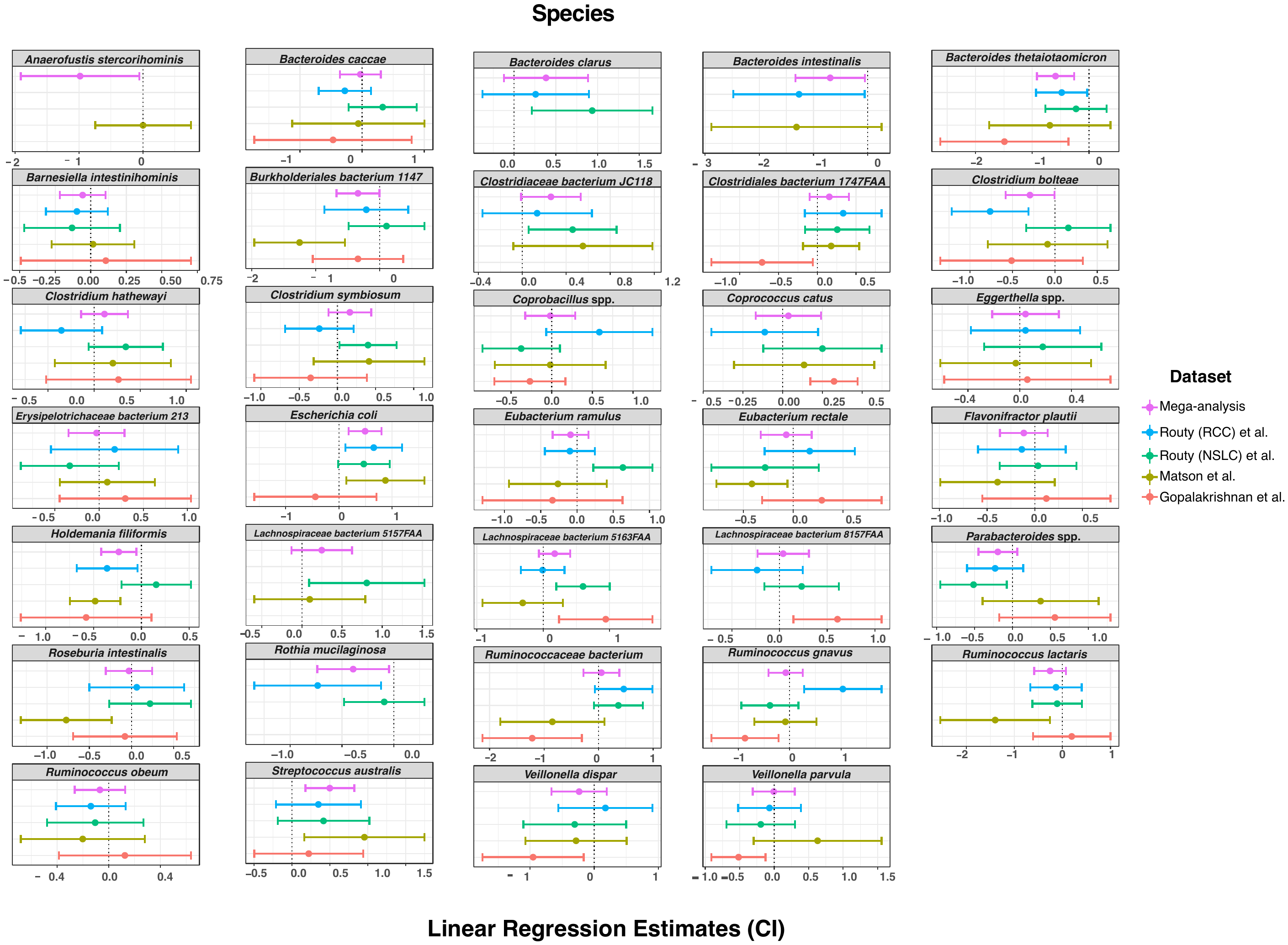

**Figure S1: Linear regression estimates (effect size) of species that are enriched or depleted in responders (R) compared to non-responders by dataset and when combined using a mega-analysis**. Taxa were included if they were significantly enriched or depleted in at least one dataset or mega- and meta-analyses using two-part log normal model. Taxa were excluded per dataset if the prevalence was <20%.
