## Supplementary material for "Mega- and meta-analyses of fecal metagenomic studies assessing response to immune checkpoint inhibitors": Figure S2

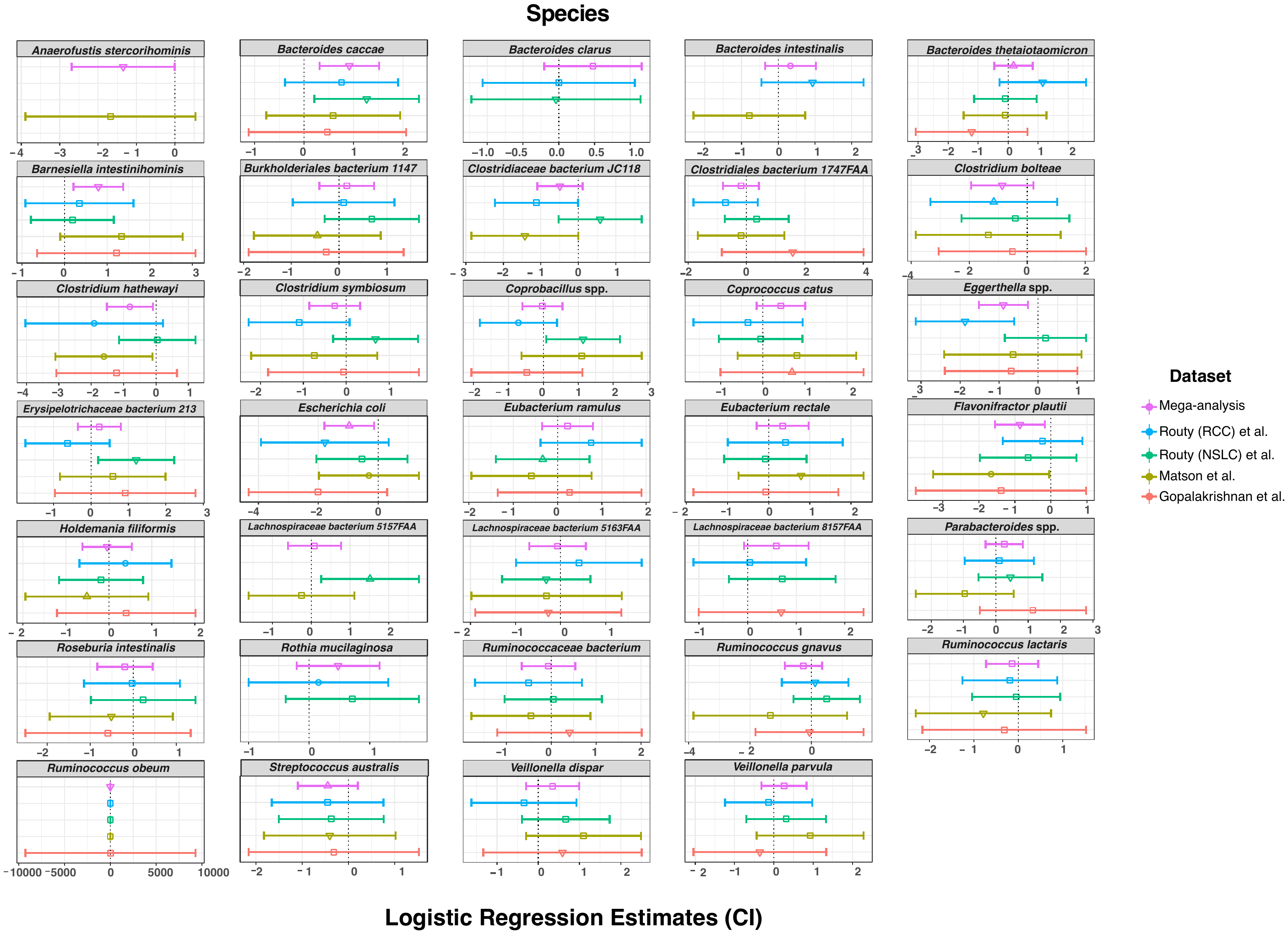

**Figure S2: Logistic regression estimates (effect size) of species based on prevalence in responders (R) compared to non-responders by dataset and combined using a mega-analysis**. Taxa were included if they were significantly different between responders and non-responders in at least one cohort or mega-meta-analyses using two-part log-normal model. Taxa were excluded per dataset if the prevalence was <20%.
