## Supplementary material for "Mega- and meta-analyses of fecal metagenomic studies assessing response to immune checkpoint inhibitors": Table S1

**Table S1:** Technical differences in sample processing and sequencing by study.

| **Study** | **Patient population** | **DNA extraction method** | **Sequencing technology** | **Metagenomic microbial taxonomic annotation** | **Read length (base pairs)** | **Average reads/ sample (million)** |
| --- | --- | --- | --- | --- | --- | --- |
| Matson *et al*. 2018 | Metastatic melanoma | QIAamp PowerFecal DNA Kit | Illumina NextSeq | MetaPhlAn 2 | 2x150 | 80.4 |
| Gopalakrishnan *et al*. 2017 | Metastatic melanoma | MO BIO PowerSoil DNA Isolation Kit | Illumina HiSeq | NCBI blast N (2016) | 2x100 | 15.9 |
| Routy *et al*. 2017 | Advanced NSCLC + RCC | Custom-DNA extraction^22,23^ | ThermoFisher Ion-Proton | NCBI blast N (2016) | 1x150 | 22.7 |

NSCLC = non-small cell lung carcinoma

RCC= Renal cell carcinoma
